# Developmental dynamics and axonal transport of cerebral cortex projection neuron mRNAs *in vivo*

**DOI:** 10.1101/2025.04.23.650263

**Authors:** Priya Veeraraghavan, Dustin E. Tillman, Melody I. Ross, Jeffrey D. Macklis

**Affiliations:** Department of Stem Cell and Regenerative Biology, and Center for Brain Science, Harvard University, Cambridge, MA, USA

**Author notes:** These authors contributed equally to this work (Priya Veeraraghavan, Dustin E. Tillman). Correspondence should be addressed to J.D.M.

## Abstract

Appropriate development of long-range nervous system circuitry requires dynamic regulation of subcellular mRNA localization, but little is known about how specific neuron subtypes control this critical process *in vivo*. Here, we employ an integrated genetic and temporally controlled approach to investigate *in vivo* developmental mRNA dynamics in somata and axons of a prototypical cerebral cortex projection neuron subtype, callosal projection neurons (CPN), which connect the cortical hemispheres to integrate sensorimotor information, and regulate high-level associative cognition and behavior. We identify cis-regulatory elements that are associated with mRNA turnover in CPN somata, including specific 3’ untranslated regions (UTRs), and elucidate distinct modes of axon transport in CPN axons for function-specific mRNA classes. Together, these findings elucidate how developing CPN control subcellular mRNA localization, and how dysregulation of these processes might lead to neurodevelopmental and/or neurodegenerative disorders. More broadly, this work identifies general mRNA localization mechanisms that likely function across projection neuron subtypes, and in other polarized cells.

## Introduction

During development, projection neurons of the cerebral cortex (in particular, the evolutionarily recent neocortex) construct specific circuits by extending long axons (hundreds to thousands of cell body (soma) diameters long in mouse; longer in developing humans) to precisely innervate distant targets^1^. Growth cones (GCs) are subcellular structures at leading edges of each axon that regulate this navigation via integration of complex extracellular cues into directional responses, before ultimately maturing into pre-synapses^2^. To effect appropriate growth and guidance, axons and GCs dynamically regulate local proteomes that are distinct from their parent somata^3,4^. This differential subcellular localization is established both by direct trafficking of proteins, and by local translation of specific mRNAs after selective stabilization, export, and transport from somata to axons and GCs^5,6^. Extremely polarized cortical projection neurons must precisely control subcellar mRNA localization, and are particularly susceptible to dysregulation of these processes, which can result in neurodevelopmental and neurodegenerative diseases^7^.

Intracellular mRNA gradients are present in neurons^8–11^, glia^12^, oocytes^13–16^, yeast^17^, and even leading edges of migrating fibroblasts^18^. Differential mRNA densities are established by a suite of RNA binding proteins (RBPs) that associate in *trans* with individual mRNAs, then direct their transport to subcellular compartments, and/or modulate their local stability^19^. RBP-mRNA interactions are driven by cis-regulatory sequences or structural elements in each individual mRNA, frequently in 3’ untranslated regions (UTRs)^20^, as well as by local concentrations of RBPs, many of which competitively bind to the same mRNA element^21^.

Several approaches enable isolation of mRNAs with cell-type and/or subcellular specificity *in vivo*. For example, translating ribosome affinity purification (TRAP) identifies actively translated mRNAs from complex tissue via genetically encoded ribosomal tags^22^. To identify mRNAs that are actively being translated, our lab recently developed experimental and analytical approaches to identify transcriptomes of subtype-specific GCs and their parent somata directly from specific neuron subtypes *in vivo* via subcellular biochemical fractionation and fluorescent small particle sorting (FSPS)^23,24^. This work has identified that GC transcriptomes are strikingly different from those of their parent somata^23^, are selectively dysregulated by deletion of key transcription factors^25,26^, are distinct across neuron sub-types and developmental stages^27^, and produce GC-localized proteins that regulate subtype-specific circuit formation^23,28^. However, mRNA subcellular localization is dynamically regulated, involving transport, stabilization, and degradation, and these approaches only enable steady-state investigation of transcriptomes and translatomes.

Dynamic regulation of mRNA stability has been studied via approaches that either induce a rapid change in transcription^29^, or introduce a non-native, chemically distinct nucleotide metabolite into newly transcribed mRNA^30–33^. Chemicals that inhibit transcription (e.g. alpha-amanitin, actinomycin D, 5,6-dichloro-1-beta-ribo-furanosyl benzimidazole (DRB)) cause widespread transcriptional arrest that alters RNA degradation kinetics^34,35^, and cannot be used *in vivo*. Temporally resolved *in vivo* metabolic labeling of neuronal projection mRNAs has only been performed in retinal ganglion cells^36^, in which partial cell-type specificity was achieved via injection into a specific tissue (e.g. the eye).

Here, we employ a combined genetic and temporally controlled approach to investigate developmental mRNA dynamics in somata and projections of callosal projection neurons (CPN), the projection neuron subtype that connects the cortical hemispheres, and plays key roles in high-level integrative, associative, cognitive, and behavioral functions^37^. We express uracil phosphoribosyltransferase (UPRT), which catalyzes the conversion of uracil to uridine, specifically in CPN, then systemically administer 4-thiouracil (4TU), leading to selective incorporation of 4-thiouridine (4sU) into CPN mRNAs. 4sU-containing mRNAs are detected via a nucleotide conversion approach^30,38^, combined with computational modeling and inference^39,40^, enabling identification of newly transcribed CPN mRNAs in both somata and axons, estimations of mRNA turnover, and functions of 3’ UTRs in regulating mRNA stability.

We identify *cis*-regulatory features in CPN mRNAs that are correlated with mRNA half-life in somata, and likely regulate transcript turnover; these include AU-rich elements, distinct RBPs, and GC localization. We also identify stabilizing/sequestering UTRs via a luciferase reporter assay, and detect distinct classes of mRNAs in CPN axons across developmental times and subcellular compartments, suggesting axonal transport mechanisms that are specific for distinct functional classes of mRNAs.

These insights into developmental mRNA dynamics of cerebral cortex projection neurons (CPN in particular) *in vivo*, combined with the experimental and analytical approaches that we develop, and partially described in dissertation work from our lab^41^, will enable future elucidation of molecular mechanisms regulating appropriate and/or aberrant mRNA stability and transport in CPN, in distinct projection neuron subtypes, and in other polarized cells.

## Results

### Time- and subtype-specific metabolic labeling *in vivo*

To identify newly synthesized transcripts in a specific projection neuron subtype with temporal resolution in vivo, we adapted metabolic labeling approaches for use in callosal projection neurons (CPN). Briefly, we introduced plasmids to CPN that encode for TdTomato and *T. gondii* uracil phosphoribosyltransferase (UPRT) via *in utero* electroporation (IUE) at embryonic day (E) 14.5 (Figures 1A-B). We then administered 4-thiouracil (4TU, 450 mg/kg) to pups at postnatal day (P) 2 or 3; 4TU is converted into 4-thiouridine (4sU) by UPRT, and incorporated into newly transcribed mRNAs^42^. We purified CPN somata with subtype specificity via fluorescence-activated cell sorting (FACS) 2, 4, 6, or 8 hours after 4TU administration, extracted RNA, and identified newly synthesized transcripts via T->C conversions, using SLAM-seq and 3 ‘end sequencing^30^. As expected, the labeling rate increases over time post-injection, and is significantly higher in UPRT-containing CPN compared to TdTomato-only CPN (Figure 1C); these control CPN display some background labeling from low activity of endogenous *M. musculus* UPRT^43^. Subcutaneous injection of 4TU led to robust labelling with minimal background, while intraperitoneal (IP) injection led to higher labeling in both background and experimental samples (Figure S1A), likely due to greater bioavailability of 4TU^44^. Therefore, subcutaneous injections were used for soma samples, and IP injections were used for lower-input axon samples (discussed below). We next investigated kinetics of individual mRNAs in CPN somata by estimating new transcript ratios (NTRs) for each transcript with a Poisson mixture model approach, using bakR^40^ (Figures 1D, S1B-E). We model relative mRNA half-lives (t_1/2_) by fitting NTRs to a model of first-order exponential decay, as previously described^39,40^. Importantly, pulse labeling of 4TU/4sU *in vivo* leads to a decrease in global labeling rate over time (Figure S1F), in contrast to constant *in vitro* labeling rates, so we scaled NTRs by a global scaling factor to account for this decrease in label (Figures S1G-I). Resulting CPN soma t_1/2_ ranged from 1.5 hours to 21.4 hours, with a mean of 7.6 hours (Figure 1E).

**Figure 1:**
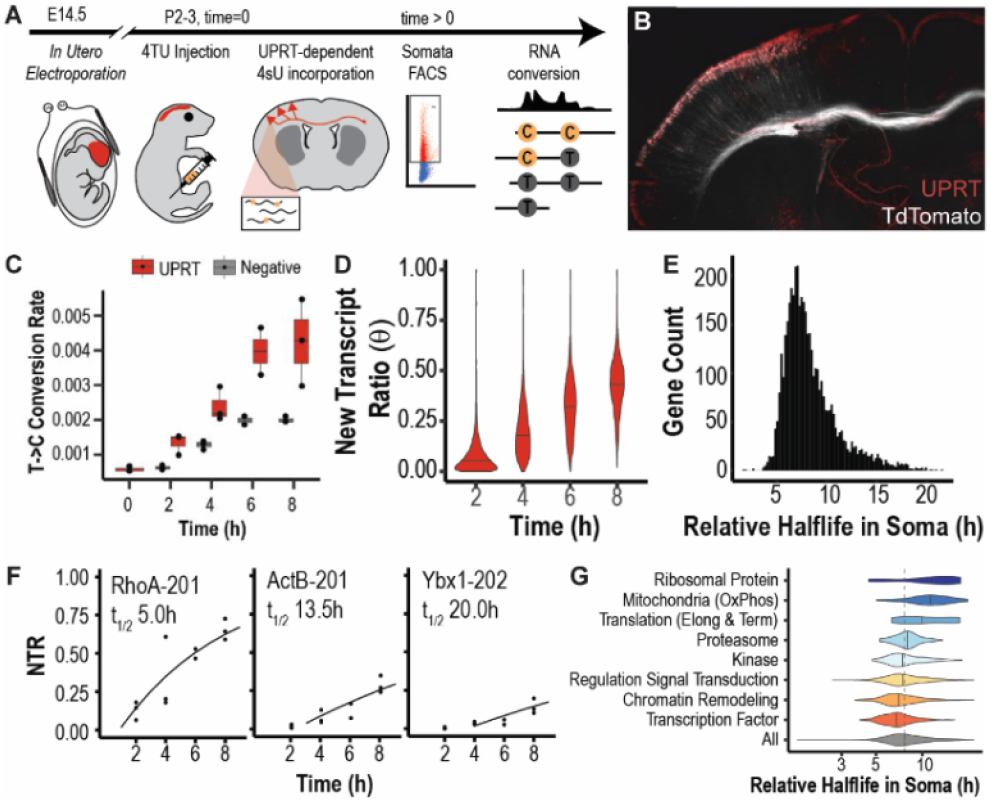
Temporally resolved, subtype-specific metabolic labeling of CPN mRNAs *in vivo*. **(A)** Schematic for metabolic labeling of CPN transcripts *in vivo*. CPN express fluorophore and UPRT via E14.5 IUE. At P2 or P3 (t = 0), 4TU is administered via intraperitoneal injection, then fluorescent CPN somata are purified at distinct time points via FACS for RNA extraction, sequencing, and conversion quantification. CPN labeled via E14.5 IUE express myristoylated TdTomato (white) and UPRT (red). **(C)** T->C conversion rate increases over time post-injection, and it is significantly higher in samples with UPRT. **(D)** Mean new transcript ratio (NTR) per gene increases over time post-injection. Distribution of transcript relative half-lives in CPN somata (RH_soma_). Changes in NTR over time, and RH_soma_ estimates for exemplar mRNAs with turnover that is fast (RhoA), medium (ActB), and slow (Ybx1). Points are per-sample maximum likelihood NTR estimates. Line is predicted curve. **(G)** RH_soma_ distributions are distinct across mRNAs associated with distinct functional gene ontology (GO) terms.

NTRs increase when new mRNA is introduced to the system (e.g. via transcription), and/or when old mRNA is removed (e.g. via degradation). We identify that per-gene transcription rates are relatively constant over the investigated time course (Figure S1J), so changes in NTR are primarily driven by removal of old transcripts. In most cells, from which the whole volume is sampled via sequencing, this removal is primarily or entirely due to degradation of old mRNAs. However, CPN, and the broad class of projection neurons, are highly polarized, so old RNA is removed from somata both by decay, and transport to subcellular compartments, particularly axons and GCs (Figure 2A). Therefore, we define the “relative half-life of a transcript” (RH) for each distinct subcompartment (e.g. RH_soma_).

**Figure 2:**
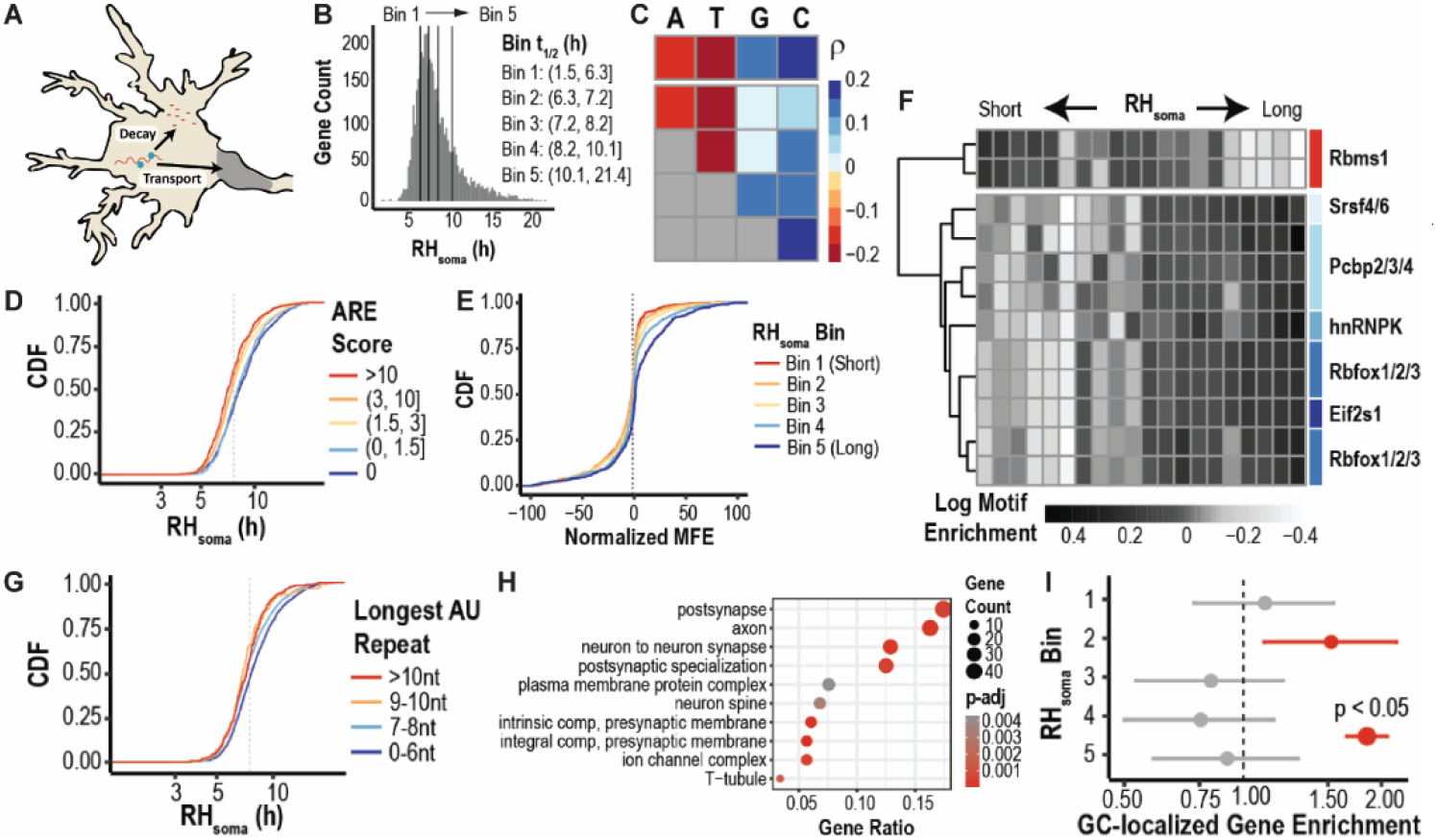
*Cis*-regulatory 3’ UTR sequence features are associated with CPN RH_soma_. **(A)** Schematic highlighting how short RH_soma_ can result from mRNA degradation and/or transport to axons/GCs. **(B)** Equal division of CPN mRNAs into five bins, based on RH_soma_. **(C)** Single- (top row) and di- (matrix) nucleotide frequencies correlate with RH_soma_. Spearman correlation coefficient. **(D)** High likelihood AU-rich elements (AREs) are negatively associated with RH_soma_. **(E)** Normalized minimum free energy of folding (MFE) is positively associated with RH_soma_. **(F)** Enrichment for RNA binding protein (RBP) motifs in 3’ untranslated regions (UTRs), based on RH_soma_. All motifs have a linear relationship between RH_soma_ and motif enrichment, with consistency between data and a fitted linear model, as measured by an F-test (adjusted p-value < 0.05). **(G)** Length of the longest AU repeat in a 3’ UTR is negatively associated with RH_soma_. **(H)** mRNAs with a longest AU repeat longer than 8 nt are enriched for synapse and axon GO terms (FDR < 0.05). **(I)** mRNAs with short RH_soma_ are GC-localized in developing CPN, as recently identified by our group^27^.

To cross-check our estimated CPN RH_soma_ values, we compared our findings to previous work in other systems^10,30,31,45–59^. Notably, per-gene RH_soma_ values are correlated across these distinct experimental contexts, indicating broad conservation of mRNA stability mechanisms, and are most highly correlated with primary cortical neurons *in vitro*^57^ (Spearman rho=0.63, Figure S1K), which most closely resemble CPN *in vivo*. Importantly, exemplar transcripts that are quickly degraded (e.g. RhoA^60^), or highly stable (e.g. Ybx1^61^), demonstrate appropriate kinetics (Figure 1F), as do gene sets defined by gene ontology (GO) terms^62,63^– mRNAs encoding for proteins that function in translation and mitochondria have longer RH_soma_ values, while Rh_soma_ values are shorter for mRNAs encoding transcription factors and chromatin remodelers^56^ (Figure 1G).

### RH_soma_ is linked with stability, degradation, and transport

Since 3’ untranslated regions (UTRs) regulate key aspects of mRNA stability and localization^20^, we next investigated whether *cis*-regulatory structures and sequence features correlate with RH_soma_ by evenly dividing transcripts into 5 bins based on RH_soma_ (Figure 2B). As predicted, UTR length only weakly correlates with RH_soma_, most notably with the slowest turnover mRNAs having the shortest UTRs (Figure S2A). This weak correlation is expected, since additional sequence elements can be either stabilizing or degrading^20^.

In contrast, 3’ UTR nucleotide content correlates with RH_soma_, with A, T, and AT-rich 3’ UTRs having shorter RH_soma_, while C and GC-rich 3’ UTRs have longer RH_soma_ (Figure 2C). To determine if this correlation is caused by AU-rich elements (AREs), known to enhance mRNA degradation^64^, we scored sequences with AREscore^65^, which assesses ARE likelihood based on the number of observed motifs and their proximity to each other. Consistent with prior work, we identify that higher AREscore values are associated with shorter RH_soma_ (Figure 2D), indicating conservation of a key mRNA degradation mechanism in developing CPN.

3’ UTRs are typically bound by RNA binding proteins (RBPs) for additional regulation of mRNA stability and localization^20^, so we next investigated whether RBP-specific sequence features correlate with RH_soma_. Intriguingly, 3’ UTR minimum free energy of folding (MFE), calculated by Vienna RNAFold^66^, then normalized for length and GC content, is lower for mRNAs with shorter RH_soma_ (Figures 2E, S2B), indicating that high-turnover transcripts have more secondary structure, and likely more RBP binding^67^.

To identify RBPs that potentially regulate mRNA stability, degradation, and/or transport, we investigated enrichment of RBP binding sites in 3’ UTRs across RH_soma_. Strikingly, mRNAs with long RH_soma_ are overrepresented for motifs bound by Pcbp and Rbfox family members, known mRNA stabilizers^68,69^, as well as other RBP motifs (Figure 2F). In contrast, Rbms1 binding sites are enriched in mRNAs with short RH_soma_, suggesting that Rbms1 might destabilize these transcripts, and/or transport them to CPN axons and GCs during development, as we recently reported^27^.

Rbms1 binds to AU repeats^70^, which localize mRNAs to neuronal projections^71^. Therefore, we investigated associations between RH_soma_ values and 3’ UTR AU repeats, and other potential indicators of axonal transport. We identify that RH_soma_ values are negatively correlated with lengths of the longest AU repeat (Figure 2G), but only 15% of these regions are likely AREs (Figure S2C), indicating that this effect is independent of ARE-mediated destabilization (Figure 2D). Further, mRNAs with long (> 8 nt) AU repeats are enriched for GO terms associated with axons and synapses (Figure 2H), and 3’ UTRs with AU repeats have greater evolutionary conservation (Figures S2D). These results indicate that AU repeats in 3’ UTRs likely target mRNAs to CPN axons and GCs in development, potentially via Rbms1. Indeed, mRNAs with short RH_soma_ values (and long AU repeats) are enriched for mRNAs that we identified as GC-localized in developing CPN during active axon elongation^27^ (Figure 2I).

### 3 ‘UTRs of Gpm6a and Gnas stabilize and sequester mRNAs

Next, we directly investigated potential functions of specific 3’ UTRs in regulating mRNA stability, degradation, and/or transport by developing and employing an *in vivo* inducible luciferase library assay. Briefly, we introduce into CPN plasmids that encode mClover and a synthetically designed library of doxycycline-dependent *R. reniformis* luciferases, each of which is fused to a distinct 3’ UTR, via E14.5 IUE (Figure 3A). We administered doxycycline (100 mg/kg) to pups at P2 or P3 via intraperitoneal injection to produce a pulse of luciferase expression, purified CPN somata via FACS 3 or 8 hours after injection, extracted RNA, and quantified UTR-specific changes in luciferase abundance via 3’ amplicon sequencing.

**Figure 3:**
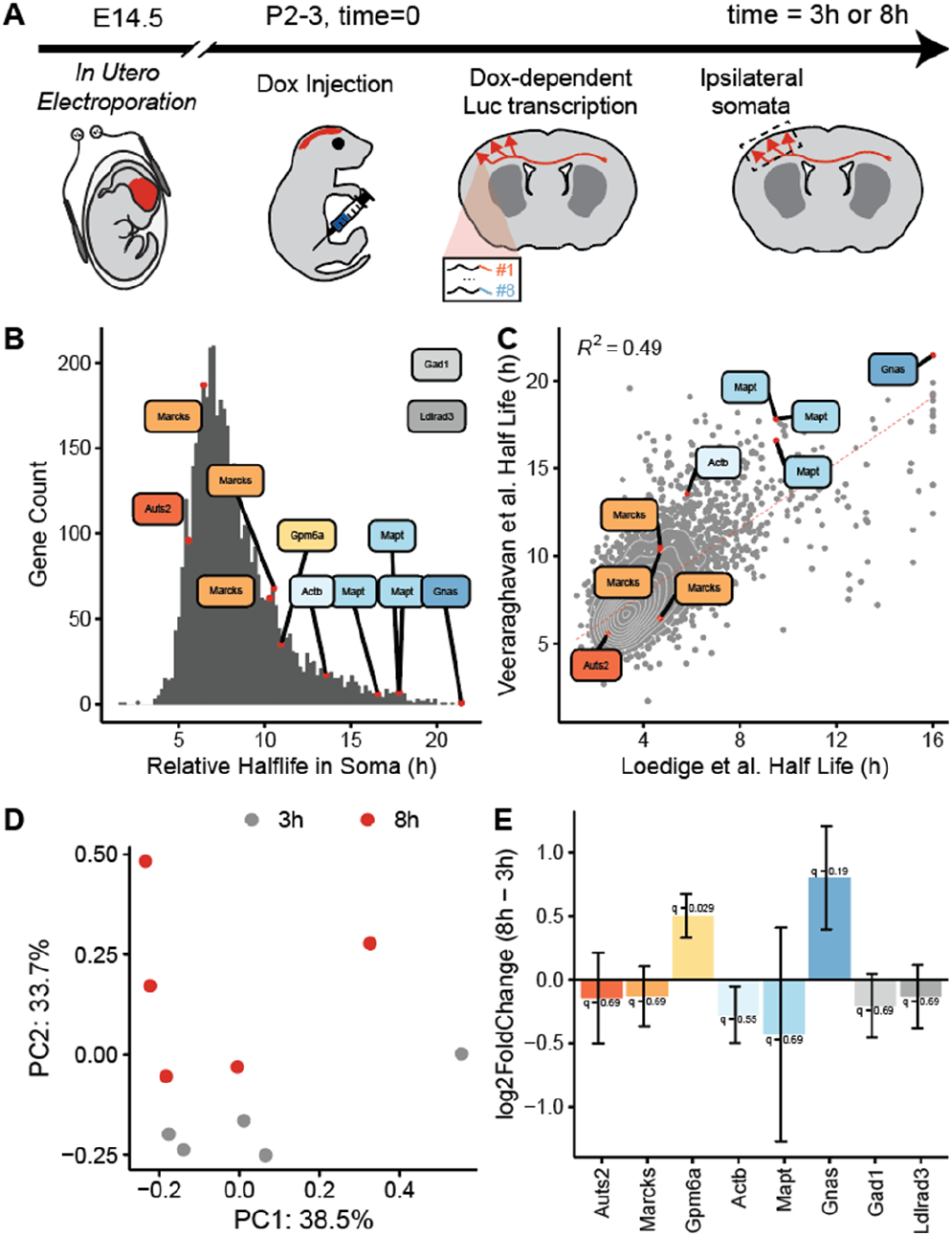
Identification of 3’ UTRs regulating mRNA dynamics in developing CPN. **(A)** Schematic for *in vivo* inducible luciferase library assay in CPN. CPN express fluorophore and a library of doxycycline-dependent luciferases, each of which is fused to a distinct 3’ UTR, via E14.5 IUE. At P2/P3 (t = 0), doxycycline is injected into intraperitoneal cavity, then CPN somata are fluorescently purified at distinct times via FACS for RNA extraction, sequencing, and differential expression analysis. **(B)** 3’ UTRs are associated with localization to cell bodies (Ldlrad3), GCs (Gad1), or span a range of RH_soma_. **(C)** 3’ UTRs have similar RH_soma_ in primary cultured cortical neurons. **(D)** Time-matched samples cluster via principal component analysis (PCA). **(E)** Luciferases fused to 3’ UTRs from Gpm6a or Gnas significantly increase over time.

To identify UTRs with distinct roles in mRNA dynamics, we investigated 3’ UTRs from Gad1 and Ldlrad3, which we recently identified as GC- and soma-localizing^27^, respectively, and 6 UTRs that span a range of RH_soma_ (Figure 3B): Auts2 (5.6 h); Marcks (6.4 – 10.5 h); Gpm6a (11.0 h); Actb (13.6 h); Mapt (16.6 –17.8 h); and Gnas (21.4 h). Importantly, these UTRs have similar RH_soma_ in primary cultured neurons from cortex^57^ (Figure 3C), increasing confidence that *in vivo* functions of these UTRs are conserved in CPN.

As predicted, all barcodes are identified with high confidence by thousands of unique molecular identifiers (UMIs; Figures S3A-B), and time-matched samples cluster via principal component analysis (PCA; Figure 3D). Importantly, library and mClover UMIs increase over time (Figure S3C), and are linearly correlated (Figure S3D), so we normalized each UTR UMI count by both factors prior to differential expression analysis via DESeq2^72^.

Strikingly, 3’ UTRs of Gpm6a and Gnas lead to a significant increase in luciferase mRNA over time (Figure 3E), suggesting that these motifs stabilize CPN mRNAs, and/or sequester them from axons and GCs. We attempted to investigate functions of 3’ UTRs in subcellular targeting by repeating this experiment with extraction of mRNA from micro-dissected CPN axons, rather than from purified CPN somata (Figure S3E). However, few UMIs were identified (Figures S3F-G), potentially due to insufficient CPN axon enrichment, bottlenecking of mRNAs in somata, and/or poor overlap between CPN labeling and doxycycline exposure. This negative result further motivated our investigations of *in vivo* CPN axon mRNA dynamics via metabolic labeling.

### Identification of mRNAs transported to CPN axons

To identify CPN mRNAs that are subcellularly localized *in vivo*, we adapted our metabolic labeling approaches for axons. Briefly, we introduced plasmids to CPN that encode for TdTomato and UPRT via E14.5 IUE, administered 4TU to pups at P2 or P3, and extracted RNA from microdissected axons 12 or 18 hours after 4TU administration that are either pre-midline (“proximal”), or post-midline (“distal”), for SLAM-Seq^30^ (Figures 4A, S4A-B). Critically, we account for variable IUE efficiencies by quantifying relative levels of reporter mRNAs fused to an ActB axonal trafficking zip-code via qPCR^73^ (Figure S4C), and by quantifying TdTomato abundance via western blot after normalization by the axonal protein Ncam^74^ (Figure S4D). We then estimate enrichment of conversions in UPRT samples with a negative binomial generalized linear model (GLM), and identify significantly labeled transcripts via a one-sided Wald test (FDR < 0.2; Figures S4E-J). As predicted, we identify transcripts in CPN axons that others have described as localizing to, and/or being locally translated in, axons and GCs (e.g. RhoA^75^, Cstb^76^, Smad5^77^; Figure 4B).

**Figure 4:**
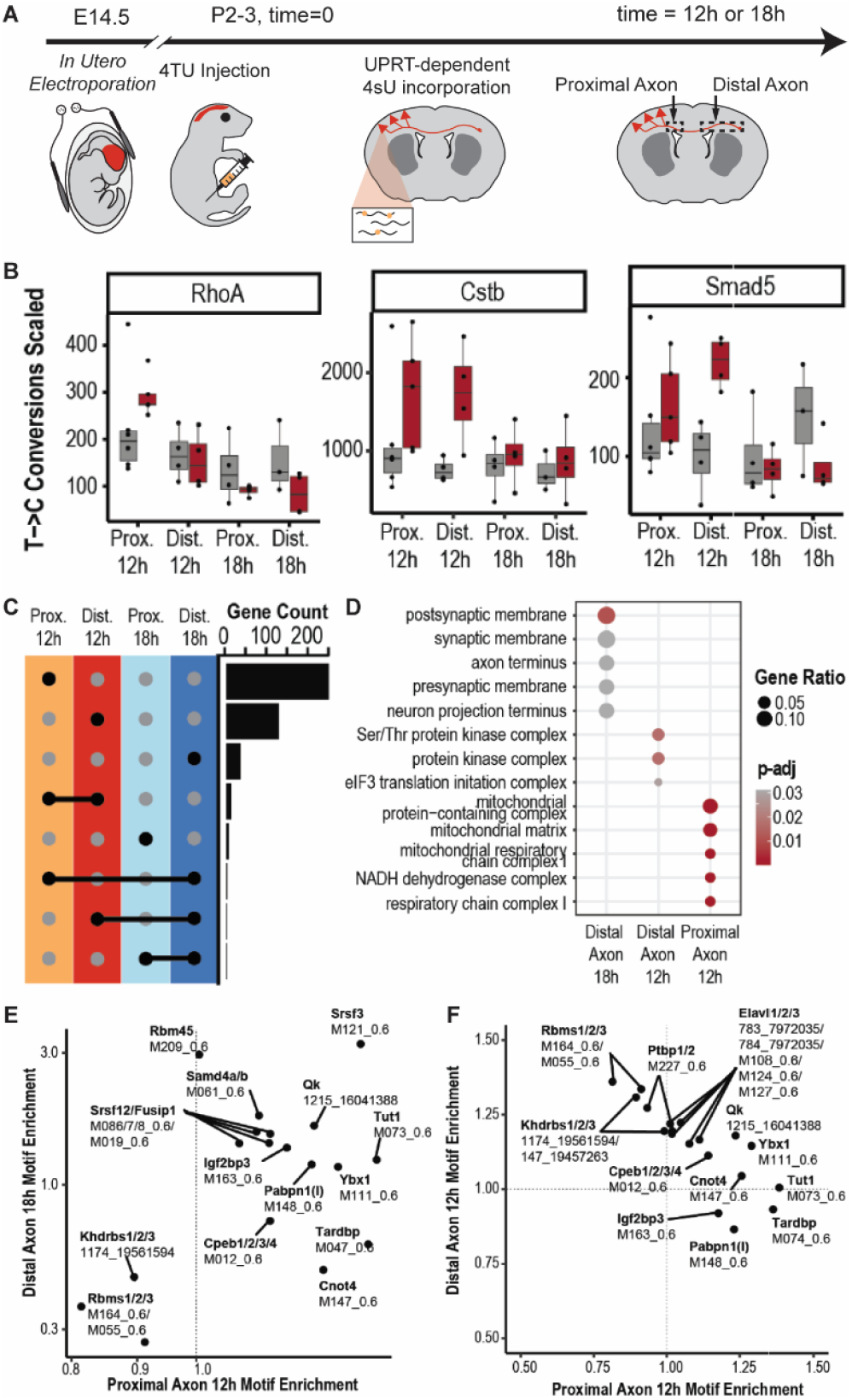
Identification of mRNAs in CPN axons across spatial and temporal domains. **(A)** Schematic for metabolic labeling of CPN axonal mRNAs *in vivo*. **(B)** Identification of exemplar genes in CPN axons that have been previously identified to localize to, and/or be locally translated in, axons (RhoA, Cstb, Smad5), as demonstrated by significant T->C conversions over background. **(C)** Labeled axonal gene sets are distinct across temporal and spatial domains. **(D)** Labeled axonal gene sets are enriched for distinct GO terms across temporal and spatial domains. **(E, F)** Labeled axonal gene sets are enriched for distinct RBP binding motifs across temporal and spatial domains.

Strikingly, labeled gene sets have minimal overlap across axon regions and time points (Figure 4C), and distinct GO terms are overrepresented in each gene set (Figure 4D), suggesting distinct mechanisms regulating function-specific mRNA transport. Further, these gene sets have differences in RH_soma_ (Figure S4K), 3 UTR length (Figure 4SL), and RBP binding sequences (Figures 4E-F). In particular, mRNAs in distal axons 12 hours after 4TU administration, the most rapidly transported mRNAs, are overrepresented for AU- and A-rich motifs that are bound by Rbms and Khdrbs RBP families, while these sequences are depleted in all other gene sets. Intriguingly, this group of mRNAs also has longer AU repeats in their 3 UTRs (Figure S4L), which are located closer to their coding sequences (CDS; Figure S4M), and include axon-localized mRNAs Cstb^76^ and Smad5^77^, as well as Efr3a, which previously has been identified to regulate CPN radial migration via Rbms1 binding^78^. Together, these results both identify *in vivo* CPN mRNAs in axons and indicate distinct RBP-mediated transcript transport mechanisms based on functional protein class.

## Discussion

### Dynamics and transport of CPN mRNAs *in vivo*

In this work, we develop an approach for subtype-specific, *in vivo* metabolic labeling of newly transcribed mRNAs with temporal resolution, enabling developmental investigation of mRNA dynamics and subcellular localization, as well as molecular mechanisms regulating these critical processes. We employ cerebral cortex callosal projection neurons (CPN) as a prototypical highly polarized neuron subtype. In particular, we quantify *in vivo* mRNA half-lives in CPN somata (RH_soma_), and validate our estimates via comparison to exemplar transcripts, previously identified dynamics of gene classes, and *in vitro* work by others (Figures 1, S1). We also identify cis-regulatory features in 3 ‘untranslated regions (UTRs) that regulate mRNA dynamics in developing CPN, including nucleotide content, motifs for distinct RNA binding proteins (RBPs), and signatures of soma export (Figures 2, S2). We identify that 3 ‘UTRs of Gpm6a and Gnas confer stability, and/or sequester mRNAs from axons, in CPN by employing an *in vivo* luciferase library reporter assay (Figures 3, S3). Finally, we adapt our subtype-specific, *in vivo* metabolic labeling approach for distinct subcellular compartments by integrating proximal and distal axon microdissection, and identify dynamic mRNA subcellular localization across temporal and spatial domains, indicating distinct, function-specific mechanisms regulating mRNA transport (Figures 4, S4).

Overall, this work demonstrates that this newly developed approach for *in vivo* metabolic labeling with subtype and subcellular specificity enables developmental identification of mRNA stability and localization, and elucidation of molecular mechanisms regulating these key processes, which are strikingly dysfunctional across a range of neurodevelopmental and neurodegenerative diseases.

### 3 UTR AU repeats regulate mRNA stability and transport

We identify that AU-rich sequences are associated with fast mRNA turnover in CPN somata, and, for a subset of genes with this motif, with efficient transport to axons. Both phenomena have been previously described, with AU-rich elements (AREs) making mRNAs more sensitive to degradation^79,80^, and AU repeats designating mRNAs for transport to axons^81^. However, little is known about how these processes are regulated by distinct RBPs.

Intriguingly, we identify that binding sites for Rbms1, a known regulator of CPN mRNA stability via AU repeat binding^78^, and a regulator of CPN circuit formation^27^, are enriched among mRNAs with short RH_soma_, and among mRNAs that are rapidly trafficked to CPN axons, suggesting that Rbms1 binding at AU repeats might regulate mRNA export from CPN somata. We attempted to test this hypothesis directly via knockdown of Rbms1 in CPN, followed by RNA-seq. However, this manipulation both increased CPN death and dysregulated CPN migration to layer II/III, preventing transcriptomic investigation. Future work would be enabled by development of new approaches for precise temporal specificity over Rbms1 manipulation to further understand its function(s) in CPN development.

Elavl RBPs, highly abundant in CPN somata^27^ and promoters of CPN subtype identity^82^, also bind AU-rich sequences, and are overrepresented among RBP binding sites in rapidly trafficked axonal mRNAs, suggesting potential function in mRNA export from CPN somata. Future work could deepen understanding of how such RBPs distinguish AREs and AU repeats from AU-rich sequences in bound RNAs for degradation and transport, respectively.

### CPN employ function-specific modes of axonal transport

We identify distinct modes of function-specific mRNA transport in CPN via the unbiased and high-throughput approaches employed here for *in vivo* metabolic labeling with temporal and subcellular resolution. For example, mitochondrial genes are overrepresented among mRNAs that slowly traffick to CPN axons, and granules with these mRNAs are enriched on Rab7a+ late endosomes, which pause in proximity to mitochondria and likely serve as translation platforms^83^, indicating a slow, organelle-based mechanism of mRNA transport. In contrast, mRNAs encoding for kinases and synaptic machinery are more rapidly transported to axons. Since rates of net anterograde axon transport are inversely proportional to granule size^83^, these mRNAs, and other mRNAs with AU-rich motifs, are likely transported at low copy numbers, possibly with even a single mRNA per granule.

### 3 ‘UTRs of Gnas and Gpm6a confer mRNA stability in CPN

We identify specific roles of distinct 3 ‘UTRs in regulating CPN mRNA dynamics *in vivo* by developing and employing a luciferase construct library. In particular, we find that the 3 ‘UTR of Gnas, which encodes for a G_s_ alpha subunit that activates intracellular signaling pathways downstream of G protein-coupled receptors (GPCRs)^84^, and that has a quite long RH_soma_, causes luciferase mRNA to be significantly stabilized and/or sequestered from axons, directly linking this sequence to CPN mRNA dynamics. Further, we identify that the 3 ‘UTR of Gpm6a, which encodes a neuronal membrane glycoprotein^85^, also stabilizes luciferase mRNA in CPN somata. We and others have demonstrated that Gpm6a mRNA is abundant in axons and neurites^23,27,57^, and that this transcript has a long half-life in neurites^57^. These results indicate that an axon-permissive signal is encoded either in full-length Gpm6a mRNA or by a factor that is introduced in *trans* co-transcriptionally.

### Rbfox proteins regulate developmental mRNA dynamics

We identify several features associated with enhanced CPN mRNA stability *in vivo*, including a lack of mRNA structure, consistent with previous reports that mRNA secondary structure leads to degradation^86^. Strikingly, binding motifs of many RBPs are enriched among low turnover mRNAs, especially those of the Rbfox family, suggesting RBP-mediated control over mRNA stability. Although Rbfox proteins are primarily known for regulating nuclear pre-mRNA splicing^87^, a cytoplasmic Rbfox1 isoform binds 3 ‘ UTRs of synaptic mRNAs associated with autism spectrum disorders (ASDs), stabilizing them by blocking miRNA binding^69^. Together, these results indicate that Rbfox1 regulates cytosolic mRNA developmental dynamics in CPN, likely via stabilization of select mRNAs.

### New approaches for *in vivo* metabolic labeling of neurons

We identify newly transcribed CPN mRNAs *in vivo* by developing and employing cell-type specific metabolic labeling approaches with temporal and subcellular resolution. Despite the success of these approaches, several opportunities for improvement remain. Boosting signal is a persistent problem *in vivo*, particularly in the central nervous system (CNS), where the blood brain barrier (BBB) blocks most systemically delivered molecules, including uracil analogs^38,88^. One possible solution is pretreatment of mice with 5-eniluracil (5EU), which inhibits the first step of uracil catabolism, potentially leading to more sustained 4TU circulation, as previously described for the chemotherapeutic agent 5-fluorouracil (5FU)^89^. Reducing background labeling is also challenging, since nucleotide scavenging pathways and endogenous UPRT paralogs prevent total noise elimination^43^. However, several groups have jointly engineered UPRT enzyme and uracil analogs to enable efficient uracil conversion with minimal endogenous scavenging. In particular, a rationally designed UPRT, combined with 5-vinyluracil (5VU), is a promising approach^33,88^.

## Conclusion

Overall, the work reported here establishes a set of experimental and analytical approaches to investigate *in vivo* mRNA dynamics of subtype-specific projection neurons with temporal and subcellular resolution. We identify developmental regulators of mRNA stability and axonal transport, enabling future studies of how these mechanisms regulate mRNA trafficking during development and in mature circuits, and how perturbation of these processes might contribute to neurodevelopmental and neurodegenerative diseases. Importantly, these approaches generalize to other neuron types, polarized cells, and/or tissues, substantially enabling and expanding investigation of *in vivo* mRNA dynamics across diverse systems and disorders.

## Acknowledgements

This work was supported by NIH Pioneer Award DP1 NS106665; Allen Distinguished Investigator Award ADI 11855 to J.D.M. from the Paul G. Allen Frontiers Group; and the Max and Anne Wien Professor of Life Sciences fund; with additional infrastructure support from NIH grants NS045523 and NS104055. P.V. was partially supported by NSF GRFP 280932, The NSF-Simons Center for Mathematical and Statistical Analysis of Biology at Harvard #1764269, and NIH Training Grant T32 GM007306-43. D.E.T. was partially supported by NIH F31 NS127518 and The NSF-Simons Center for Mathematical and Statistical Analysis of Biology at Harvard #1764269. We thank Joyce LaVecchio and Nema Kheradmand of the HSCRB-HSCI Flow Cytometry Core; the Bauer RNA sequencing core and the Harvard Center for Biological Imaging for infrastructure and support; Tim Sackton for statistical advice; and other members of the Macklis lab for helpful suggestions.

## Author contributions

J.D.M. and P.V. conceived of the project; J.D.M., P.V., and D.E.T. designed experiments; P.V. and M.R. generated constructs; P.V. performed all mouse work. P.V. and D.E.T. performed soma preps and sorts; P.V. and D.E.T. performed library preparation for RNA sequencing; D.E.T. performed western blot validation; P.V. performed histology; P.V. performed bioinformatic analysis with input from D.E.T; P.V., D.E.T., and J.D.M interpreted data and wrote the manuscript. All authors read and approved the manuscript.

## Materials and Methods

### Mice

Experiments involving mice were performed with approval of the Harvard University Institutional Animal Care and Use Committee (IACUC) and in accordance with all federal and state guidelines. All data on callosal projection neurons (CPN) were collected from wild-type CD1 (Charles River Laboratories; RRID:IMSR_CRL:022) mouse pups of both sexes via subtype-specific expression of fluorescent proteins using unilateral *in utero* electroporation (IUE) at embryonic day E14.5, targeting CPN in cortical layers II/III. Due to the exploratory nature of the investigations described here, a formal power analysis was not required or performed.

#### In utero electroporation

Subtype-specific labeling of CPN was achieved using unilateral *in utero* electroporation of genetic constructs at E14.5, as we and others have previously described^90^. Briefly, a pregnant female was anesthetized using 1.5-2% isoflurane, then prepared for surgery by applying eye ointment (#2444062, Systane), and administering buprenorphine (#60969, Par Pharmaceutical). Lower abdominal hair was removed using hair removal cream, and skin was locally disinfected with ethanol. An incision was made through skin and abdominal musculature, and the uterine horns were exposed, gently externalized from the intraperitoneal cavity onto a sterile gauze pad on the abdomen, where they were continuously kept moist using sterile, pre-warmed PBS. Embryos were individually positioned and injected unilaterally into a lateral ventricle with 0.7-1 ul DNA (5 ug/ul) supplemented with Fastgreen (#AAA16520-06, Thermo Scientific Chemicals) for visualization, using a pulled and beveled glass micropipette (#22-260-943, Fisher). DNA was electroporated into neural progenitors lining the ventricles by applying voltage across the injected cortical area (5 pulses, 50 ms on, 950 ms off, 34 V), targeting lateral sensorimotor areas of the cortical plate. Following electroporation, embryos were gently placed back into the intraperitoneal cavity, and the wound was closed using sutures (#39010, Covetrus). The pregnant dam was allowed to fully recover from anesthesia and surgery on a thermally-controlled mat under a low-intensity heat lamp before being returned to animal husbandry. Postoperative care following surgery included daily checks of overall well-being and the wound, and administration of buprenorphine (#60969, Par Pharmaceutical) every 12 hours for the first 3 days post-surgery. After birth, pups were screened for unilateral cortical fluorescence using a fluorescence stereoscope, and successfully electroporated pups were marked with a tail clip and included in downstream experiments, while unsuccessfully electroporated pups were excluded from downstream experiments - Attrition rate of electroporation was approximately 30% across all experiments.

### DNA constructs

Constructs were derived from pCAG-[MCS]-IRES-GFP (CBIG), courtesy of C. Lois. For metabolic labeling, TdTomato (RRID:Addgene_26771) and UPRT (RRID:Addgene_47110) were cloned downstream of pCAG to create pCAG-myr-TdTomato and pCAG-myr-TdTomato-2A-UPRT. For *in vivo* luciferase library experiments, mClover (RRID:Addgene_74236) and luciferase (RRID:Addgene_38235) were cloned downstream of pCAG to create pCAG-mClover and pCAG-luciferase. Luciferase was then fused downstream of distinct 3’ UTRs via InFusion cloning of custom primers (Azenta Life Science). All constructs were prepared using the Endotoxin-free Maxiprep kit (Qiagen, #12362), following manufacturer’s instructions, and quality was verified using restriction enzyme digests and Sanger sequencing (Azenta Life Science) prior to CPN labeling via E14.5 IUE.

### 4-thiouracil injection

P2 or P3 pups were cryoanesthetized on ice for 60 seconds each, then 4-thiouracil (Sigma) was diluted in DMSO to 400 mg/kg, and injected subcutaneously into fat pads on back of neck, or into the intraperitoneal cavity, using a 30cc insulin syringe. Pups fully recovered on a thermal mat.

### Doxycycline injection

P2 or P3 pups were cryoanesthetized on ice for 60 seconds each, then 20 ul (10 mg/uL) of doxycycline diluted in DMSO was injected into intraperitoneal cavity via Hamilton syringe. Pups fully recovered on a thermal mat

### Subtype-specific soma isolation and purification via FACS

All metabolic labeling experiments were performed under red light. Soma isolation and subtype-specific purification via FACS were performed as we and others have previously described (https://doi.org/10.1038/s41596-021-00638-7)^24,91^. Briefly, P3 pups, which had been electroporated at E14.5 to label CPN, were deeply anesthetized in ice and decapitated, enabling brain extraction from the skull. Fluorescently labeled neurons in layer II/III were microdissected in ice-cold HBSS (Thermo 14025134) via a fluorescence stereomicroscope, and transferred to a Falcon tube with dissociation solution (DS). Microdissected cortical tissue containing CPN from 2-3 pups was used for each sample. Tissue was washed twice with 3-5 mL DS, then enzymatically dissociated by incubating twice with 5 mL enzyme solution (ES) supplemented with DNaseI for 15 min per incubation. ES was removed, and tissue was washed twice with wash solution (WS), then mechanically dissociated via gentle trituration with a fire polished glass Pasteur pipet. Dissociated CPN were washed in 10 mL WS and pelleted at 80*g* for 5 min at 4 °C. Supernatant was removed, and neurons were resuspended in 500 ul WS using a fire polished glass Pasteur pipet. Subtype-specific, labeled CPN were sorted into RLT buffer (Qiagen) supplemented with 10% beta mercaptoethanol (Thermo 31350010) by blinded machine operators using a customized SORP FACSAriaII (BD Instruments) equipped with a 300 MW 488 nm laser set to 100 MW with large beam height, and an 85 um nozzle run at 45 psi, with the UV laser turned off, to prevent potential metabolic labeling interference.

### Axon microdissection

All metabolic labeling experiments were performed under red light. P3 pups, which had been electroporated at E14.5 to label CPN, were deeply anesthetized in ice and decapitated, enabling brain extraction from the skull. Fluorescent axons from somata in layer II/III were microdissected from the corpus callosum in ice-cold HBSS (Thermo 14025134) using a fluorescence stereomicroscope. “Proximal axons” were dissected from the ipsilateral hemisphere using both a lateral boundary to avoid the electroporated somata and a medial boundary located 2 mm from the midline. “Distal axons” were dissected from the contralateral hemisphere from 2 mm from the midline to the edges of visible fluorescence. Microdissected regions from 2 pups were used for each sample. Axon segments were placed in 350 uL RLT (Qiagen AllPrep Kit) supplemented with beta mercaptoethanol (Thermo 31350010). Tissue was homogenized with a 26G needle and syringe prior to RNA isolation.

### RNA isolation

For metabolic labeling experiments, RNA was isolated via a Qiagen AllPrep Kit, according to manufacturer’s instructions, using RNeasy MinElute columns and the following modifications: buffer RPE was supplemented with beta mercaptoethanol (Thermo 31350010) to a final concentration of 10 mM, and RNA was eluted in 1 mM DTT (Thermo P2325). For metabolically labeled axon samples, protein was extracted from the initial flow-through, according to manufacturer’s instructions, solubilized in 5% SDS, and quantified using the Qubit broad range protein assay for downstream normalization analysis. RNA was carboxyamidomethylated in a 50 ul reaction, as previously described^92^ (50% DMSO (Sigma 41640), 10mM iodoacetamide (Sigma I1149), 50mM sodium phosphate buffer pH 8.0 (Sigma 74092 and 94046), and 1x Superase RNase Inhibitor (Invitrogen AM2696)) at 50 °C for 15 min, then quenched with 1 ul of 1 M DTT (Thermo P2325). RNA cleanup was performed with Zymo Clean and Concentrator-5. RNA was eluted in 10 ul water, and samples were stored at −80 °C. Otherwise, RNA was isolated via a Qiagen AllPrep Kit using RNeasy MinElute columns. RNA quality was assessed on a Tapestation 2100 with the High Sensitivity RNA kit (Agilent) prior to library preparation and RNA sequencing.

#### Library preparation and RNA sequencing

For metabolically labeled somata, the QuantSeq FWD kit with hexamer UMIs (Lexogen) was used for library preparation, using ~20-100 ng RNA per sample, then a NovaSeq (Ilumina) was used for 100 bp single-end reads via 3’ end sequencing for ~25-40M reads per sample. For axons, libraries were prepared with SmartSeq v4 (Takara Bio) ¼-volume reactions, using ~5 ng RNA per sample, then a NovaSeq (Ilumina) was used for 2×150 bp paired-end reads via full-transcript sequencing (increasing coverage and detection of T->C conversions) for ~25-40M reads per sample. For luciferase library experiments, RNA was reverse transcribed with oligo-dT primers and the SuperScript IV First-Strand Synthesis System (Invitrogen 18091050), then primers with overhangs compatible with distinct 3’ UTRs and Ilumina indexing primers were used to amplify desired mRNA regions. qPCR was performed with Lexogen PCR Add-on and Reamplification Kit V2 (Lexogen 208.96) to determine optimal cycle numbers, then indices were added with Lexogen UDI 12 nt Unique Dual Indexing V2 Add-on Kit B1 (Lexogen 202.96). A NextSeq (Ilumina) was used for 2×50 bp paired-end reads via full-transcript sequencing for ~4M reads per sample.

### Perfusion, tissue processing, and immunocytochemistry

Postnatal mice were deeply anesthetized in ice and transcardially perfused with 3-5 mL of ice cold PBS, followed by 5 mL of cold 4% paraformaldehyde (PFA). Brains were extracted from the skull, and postfixed in 4% PFA overnight. The next day, perfused brains were washed in PBS, and cryoprotected with 30% sucrose in PBS. Tissue was embedded in O.C.T. (#25608-930, Sakura Finetek USA Inc), sectioned via cryostat (Leica, 50 um coronal sections), and collected in PBS with 0.025% azide. For immunocytochemical labeling, tissue was blocked in PBS with 0.3% TritonX100, 0.3% BSA, and 2% donkey serum for 1 hour at room temperature, then incubated with primary antibodies (RFP #600-401-379 Rockland; HA #H3663 Sigma) in blocking buffer on a rocker overnight at 4 °C. The next day, sections were washed 3 times in PBS on an orbital shaker, incubated for 2-3 hours at room temperature with fluorescently-conjugated secondary antibodies (Life Technologies) and DAPI (1:10,000, #D1306 Thermo), washed 3 times with PBS, mounted on glass slides (Supefrost, #48311-702 VWR), coverslipped with fluoromount (#0100-01 VWR/Southern Biotech), and sealed with nail polish. Images were acquired via an epifluorescence microscope with a motorized stage and a 10x objective (Nikon NiE). Whole sections were imaged via EDF z-stack projections and mosaic image stitching with NIS Elements (Nikon).

### Sequence alignment and conversion calling

Sequences were aligned to GRCm38^93^ via Next-Gen Map^94^ with the “-slam” option to account for T->C nucleotide conversions. SNPs were called for each sample, using samtools mpileup (RRID:SCR_002105) and VarScan2^95^, and all high-quality variants were pooled to generate a reference for SNPs in our CD1 mouse strain (RRID:IMSR_CRL:022). Custom scripts, publicly available on GitHub, were used to detect conversions in the read sequence compared to the reference while masking SNPs.

### Estimating new transcript ratios

NTR was estimated as others have previously described^40^, with some slight modifications. Briefly, the conversion error rate (p_old_) was calculated as the sample conversion rate of unlabeled samples. In each labeled sample, the likelihood of observing a certain number of T->C conversions in a read was parameterized by a Poisson mixture model:

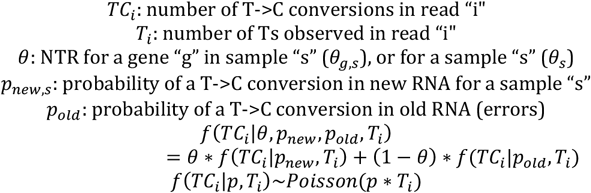

For sample-wide estimates, the maximum likelihood estimates for p_new,s_ and for θ_s_ were obtained. The p_new,s_ was then used to calculate maximum likelihood estimates for θ_g,s_ for each gene in every sample

### Estimating half-lives

Relative half-lives were determined as follows:

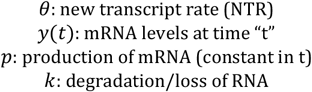

θ estimates follow from the equation describing steady-state mRNA levels:

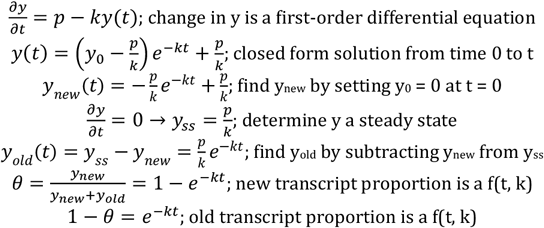

The estimate for half-life (t_h_) then follows:

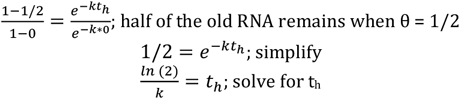

### Identification of labeled transcripts in axons

Since axonal transcripts are expected to be mature mRNAs, we accounted for variability across mouse 4TU injections by normalizing conversions per gene based on the median intron conversion rate across each sample (Figures S4E-F). We set a gene detection threshold of > 50 reads and > 1.5 normalized conversions per 1,000 Ts, which was required in >2/3 of samples of a particular sample type to include that gene for additional downstream analyses (Figure S4G). A negative binomial adaptation of a GLM was then fit to these data (using glm.nb from the MASS package in R). The default log link function was used for the response variable TC_scale:g,s_, which we defined as follows:

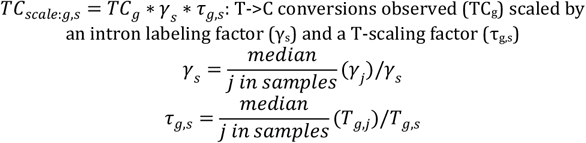

And the linear term is:

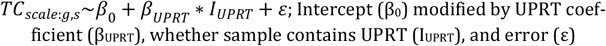

The model was first fit with all data, then (1) with “leave-one-out” cross validation (remove 1 sample in each training run, then assess model performance on the held-out sample), and (2) with “bootstrapping” (randomization of group assignments to simulate a null distribution of β_UPRT_ estimates). Significance was assessed using a one-sided Wald test for β_UPRT_ coefficient, and p-values were corrected via Benjamini-Hochberg procedure.

### RNA features analysis

#### UTR Sequences

QuantSeq soma reads were aligned to GRCm38. To define associations between 3’ UTRs and observed reads, a pileup of 5’ ends was generated, then intersected with 3’ UTRs from Ensembl version 101 (RRID:SCR_002344). If a pileup was within 250bp of a 3’ end, it was assigned to that 3’ UTR isoform. If a pileup was further from a 3’ end (typically > 500bp), a new “truncated” 3’ end isoform was called. 3’ UTR sequences were retrieved from Ensembl bioMart, and “truncated” isoform sequences were generated as a substring of annotated parent 3’ UTRs. These UTR sequences were applied to axons.

#### Normalized minimum free energy of folding (MFE)

All sequences were run through Vienna RNAFold with standard paramaters to calculate MFE. Simulated background sequences were randomly generated with varying %GC content (20-80%) and length (50-10,000 bp). Each “true” sequence was binned by %GC content and length, and the respective background was used to calculate normalized MFE, as previously described^95^: 100*(MFE_observed_-MFE_simulated_)/(L-L_0_); L_0_ = 8.

#### AU-rich element (ARE) score

For each sequence, ARE scores were calculated as previously described^65^. For Figure 2D, whole 3’ UTR sequences were used to calculate ARE scores without length normalization. For Figure S2C, ARE scores were computed using 50 bp windows to enable comparisons between all 3’ UTR regions and all AU repeat centered regions. Motif enrichment: Enrichment for RBP motifs was computed using the Transite program and R package^97^, which employs a hypergeometric test to compare motif prevalence in a subset relative to background.

#### Motif scanning

FIMO^98^ from the MEME suite was used to scan sequences for particular motifs. Standard parameters were used.

#### Evolutionary Conservation Analysis

phyloP^99^ scores for placental mammals were obtained from the UCSC Genome Browser (RRID:SCR_005780), and average scores across a 50 bp genomic window were computed using bigWigAverageOverBed^100^. Windows were centered on AU repeats, or tiled across entire 3’ UTRs with 10 bp steps.

**Figure S1:**
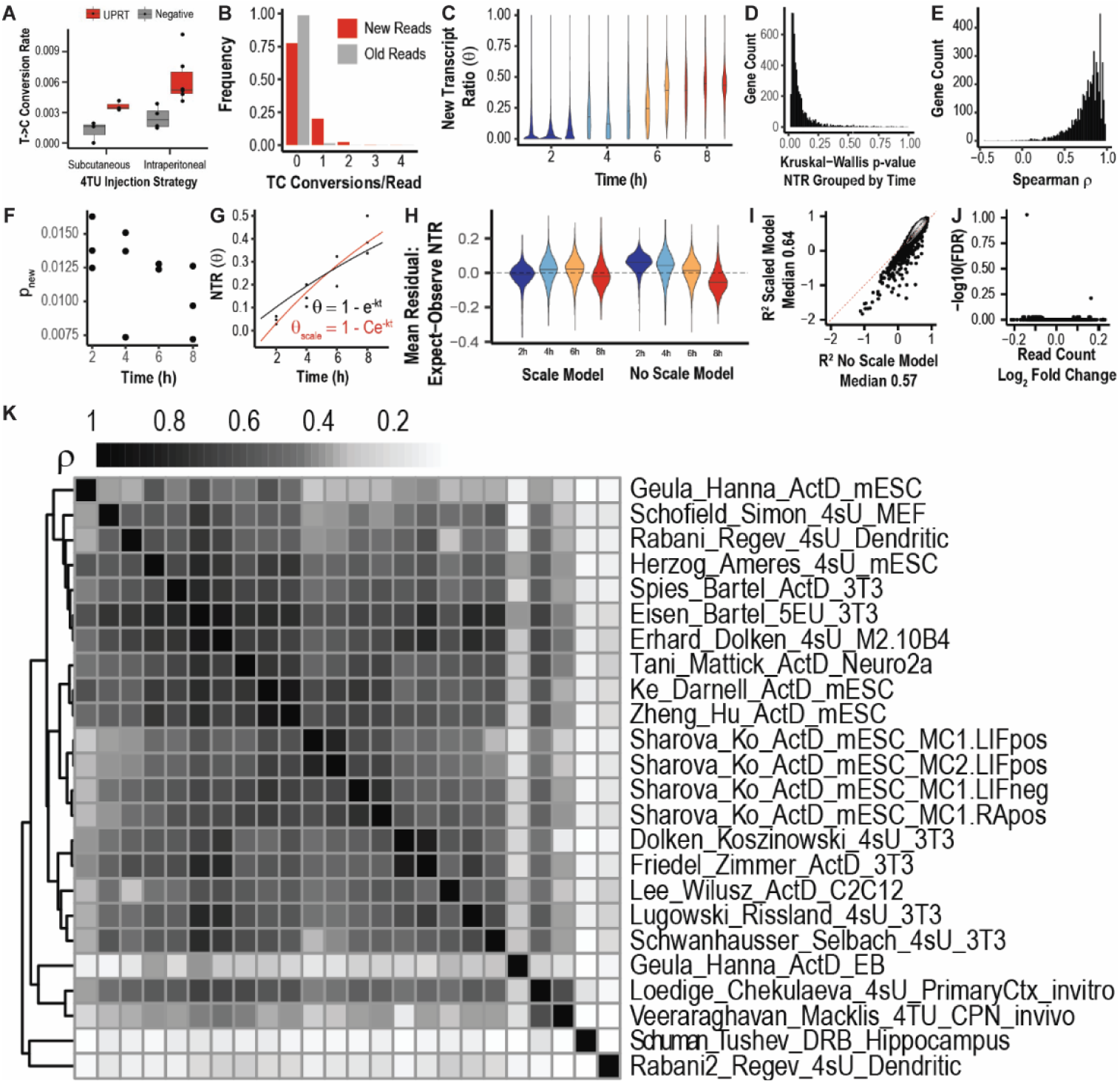
Identification of CPN transcript half-lives via *in vivo* metabolic labeling. **(A)** Intraperitoneal injection of 4-thiouracil (4TU) leads to more significant incorporation over background. **(B)** New reads are more likely to have T->C conversions (true label) than old reads (background label). (c) Genome-wide, per-sample new transcript ratio (NTR) estimates increase after 4TU injection. **(D)** Per-gene comparisons of NTR within a timepoint group versus between timepoint groups identifies that most variability arises from differences between timepoints, rather than from within timepoints. **(E)** Per-gene Spearman correlation coefficient between NTR and time identifies that NTR increases over time. **(F)** Probability of T->C conversions in new RNA (p_new_) decreases over time, consistent with 4TU clearance. **(G)** Estimating k_deg_ to calculate RH_soma_ (ln(2)/k_deg_) with (red) or without (black) a scalar to account for 4TU clearance. **(H)** Residuals for the “scaled” model do not negatively correlate with time, unlike residuals for the “unscaled” model. **(I)** The “scaled” model has slightly better R^2^ goodness of fit compared to the “unscaled” model. **(J)** No mRNAs change significantly over time (q < 0.05), validating mRNA steady-state assumptions. **(K)** Among all studies, RH_soma_ Spearman correlation is best with half-lives from primary cultured cortical neurons.

**Figure S2:**
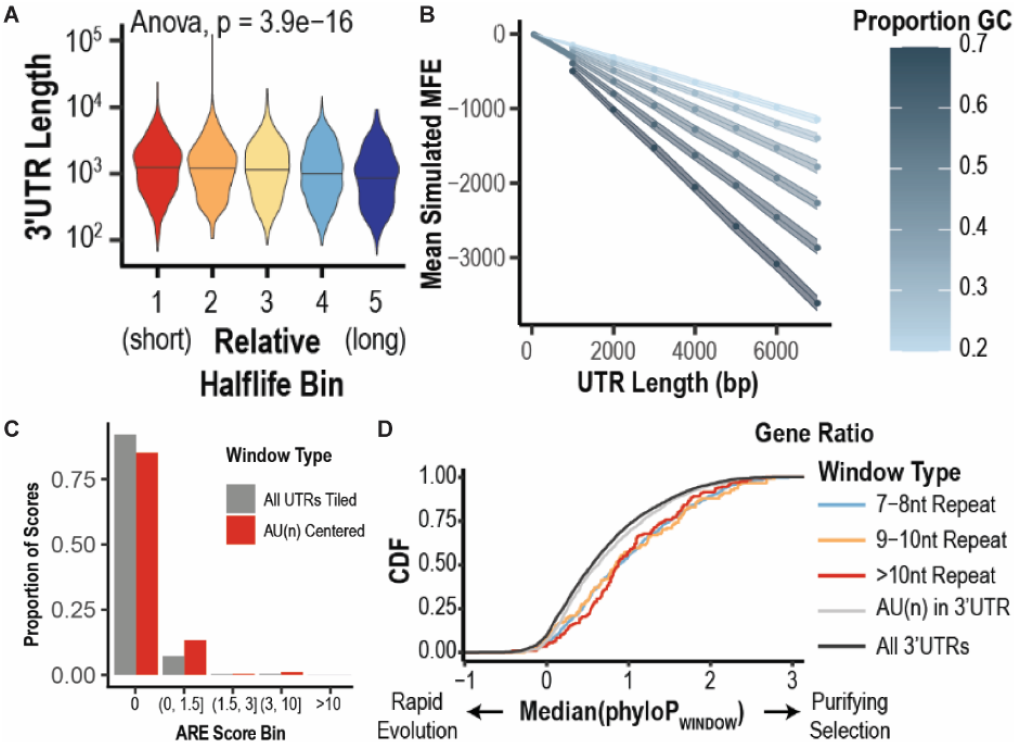
*Cis*-regulatory features are associated with *in vivo* CPN transcript half-lives. **(A)** 3’ UTR length negatively correlates with CPN soma mRNA half-life (RH_soma_). **(B)** Simulated background scores for minimum free energy of folding (MFE) vary by %GC content and 3’ UTR length. **(C)** Most AU repeats do not overlap with AU-rich element (ARE) motifs across 50 bp windows. **(D)** Per-gene AU repeats in 3’ UTRs have more evolutionary conservation.

**Figure S3:**
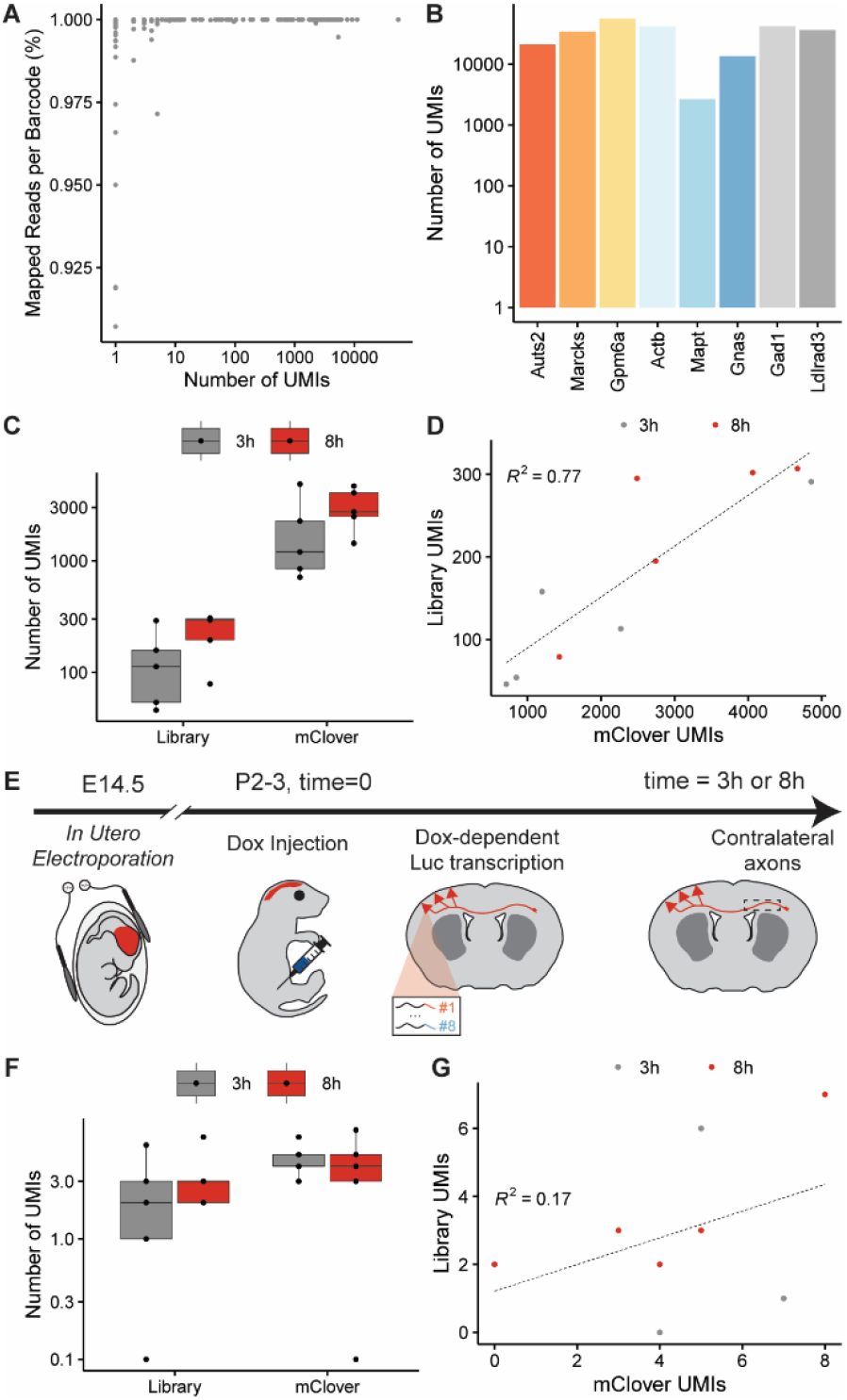
Identification of 3’ UTR barcodes from luciferase library in somata, but not axons. **(A)** All 3’ UTR barcodes are identified confidently by unique molecular identifiers (UMIs). **(B)** All 3’ UTR barcodes are identified at high abundances, as quantified by number of UMIs per 3’ UTR barcode. **(C)** UMIs for barcodes and mClover increase over time after doxycycline injection, motivating normalization by library and mClover abundances. **(D)** UMIs for barcodes and mClover are well-correlated, enabling use of both factors for normalization. **(E)** Schematic for *in vivo* inducible luciferase library assay in CPN axons. CPN express fluorophore and a library of doxycycline-dependent luciferases, each of which is fused to a distinct 3’ UTR, via E14.5 IUE. At P2 or P3 (t = 0), doxycycline is administered via intraperitoneal injection, then fluorescent CPN axons are isolated at distinct time points via microdissection for RNA extraction, sequencing, and differential expression analysis. **(F)** UMIs for barcodes and mClover are not abundant in CPN axon samples, likely due to the limited level of enrichment. **(G)** UMIs for barcodes and mClover are poorly correlated, due to low abundance of these mRNAs in CPN axons.

**Figure S4:**
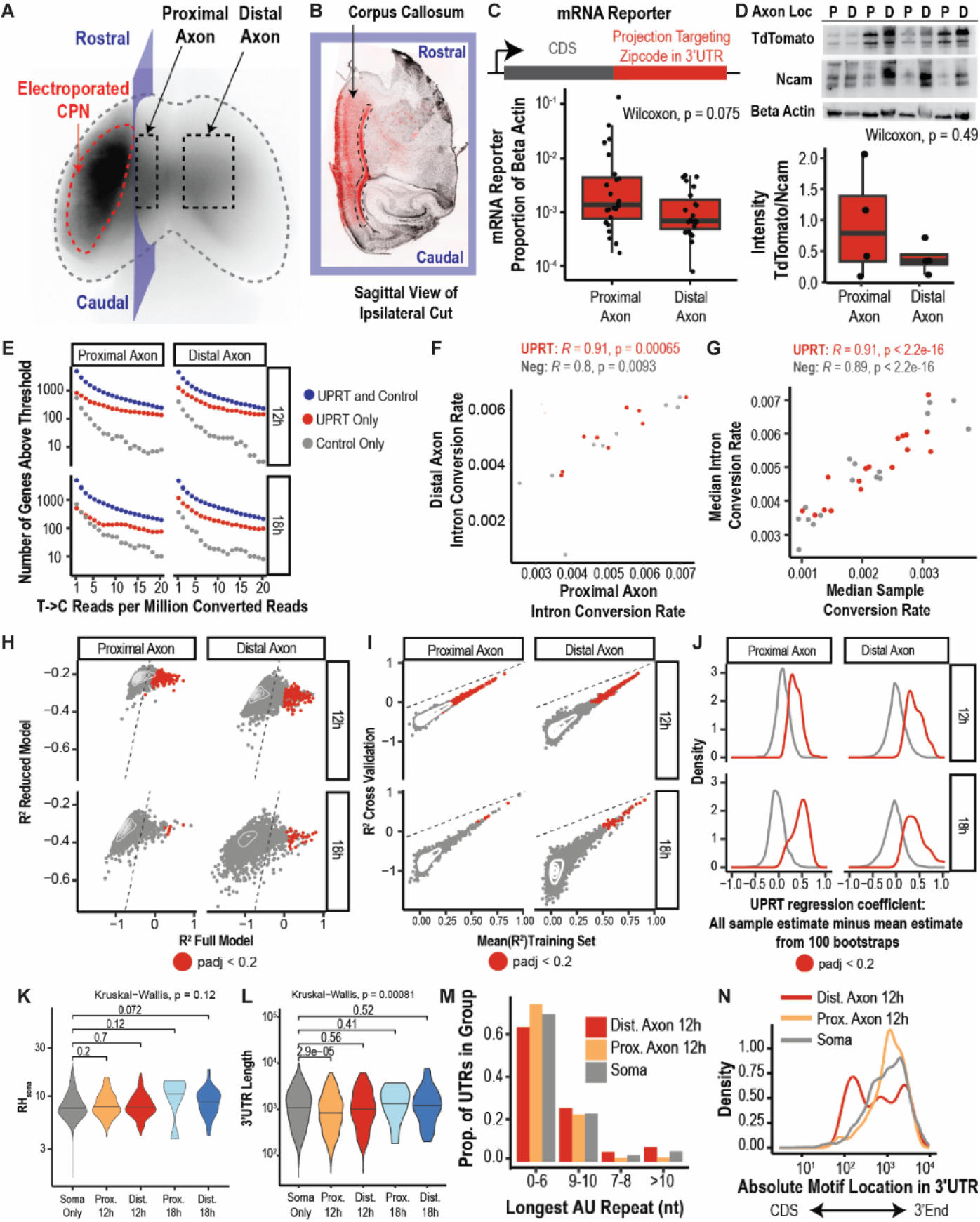
Identification of mRNAs in CPN axons via *in vivo* metabolic labeling. **(A)** CPN labeled via E14.5 IUE extend axons to contralateral cortex, as demonstrated by whole-mount imaging. **(B)** CPN axons are fluorescently microdissected, shown in the parasagittal section of the labeled brain from (A). **(C)** mRNA reporter is significantly more abundant in proximal axons (relative to beta actin), as identified by qPCR. **(D)** TdTomato appears non-significantly more abundant in proximal axons (relative to Ncam), as identified by western blot. **(E)** UPRT samples have more mRNAs above T->C conversion thresholds in ¾ of samples than non-UPRT samples. **(F)** Median intron conversion rates are well correlated between proximal and distal axons, enabling normalization. **(G)** Median intron conversion rates are well correlated with median T->C conversion rates, enabling normalization. **(H)** A negative binomial generalized linear model (GLM) performs better with a β_UPRT_ term (“full” model) than without it (“reduced” model) for some mRNAs, enabling identification of axon mRNAs. R^2^ measures “goodness of fit”. **(I)** R^2^ between “training” (one held-out sample) and cross-validation (containing the held-out sample) is positively correlated across training runs, suggesting that this model is generalizable. **(J)** “True” value of β_UPRT_ (full model – mean(bootstrapped β_UPRT_)) is strongly positive for labeled mRNAs in axons. **(K)** RH_soma_ does not change significantly across axon samples, but appears non-significantly higher in 18 hour axons. **(L)** 3’ UTR lengths change significantly across axon samples, and are significantly lower in proximal, 12 hour axons. **(M)** mRNAs in distal, 12 hour axons have longer AU repeat motifs in their 3’ UTRs. **(N)** AU repeats in mRNAs from distal, 12 hour axons are closer to coding sequences (CDS).

